# Personalized modeling of the human gut microbiome reveals distinct bile acid deconjugation and biotransformation potential in healthy and IBD individuals

**DOI:** 10.1101/229138

**Authors:** Almut Heinken, Dmitry A. Ravcheev, Federico Baldini, Laurent Heirendt, Ronan M.T. Fleming, Ines Thiele

**Affiliations:** Luxembourg Centre for Systems Biomedicine, University of Luxembourg, Esch-sur-Alzette, Luxembourg

## Abstract

The human gut microbiome performs important functions human health and disease. Intestinal microbes are capable of deconjugation and biotransformation of human primary bile acids to secondary bile acids. Alterations of the bile acid pool as a result of microbial dysbiosis have been linked to multifactorial diseases, such as inflammatory bowel disease (IBD).

Constraint-based modeling is a powerful approach for the mechanistic, systems-level analysis of metabolic interactions in microbial communities. Recently, we constructed a resource of 773 curated genome-scale reconstructions of human gut microbes, AGORA. Here, we performed a comparative genomic analysis of bile acid deconjugation and biotransformation pathways in 693 human gut microbial genomes to expand these AGORA reconstructions accordingly (available at http://vmh.life).

To elucidate the metabolic potential of individual microbiomes, publicly available metagenomic data from a cohort of healthy Western individuals, as well as two cohorts of IBD patients and healthy controls, were mapped onto the reference set of AGORA genomes. We constructed for each individual a large-scale personalized microbial community model that take strain-level abundances into account. Using flux balance analysis, we found that distinct potential to deconjugate and tranform primary bile acids between the gut microbiomes of healthy individuals. Moreover, the microbiomes of pediatric IBD patients were significantly depleted in their bile acid production potential compared with controls. The contributions of each strain to overall bile acid production potential across individuals were found to be distinct between IBD patients and controls. IBD microbiomes were depleted in contributions of Bacteroidetes strains but enriched in contributions of Proteobacteria. Finally, bottlenecks limiting secondary bile acid production potential were identified in each microbiome model. For ursodeoxycholate, the abundance of strains producing the precursor rather than of strains directly producing this secondary bile acid was synthesis-limiting in certain microbiomes.

In summary, we integrated for the first-time metagenomics data with large-scale personalized metabolic modeling of microbial communities. We provided mechanistic insight into the link between dysbiosis and metabolic potential in IBD microbiomes. This large-scale modeling approach provides a novel way of analyzing metagenomics data to accelerate our understanding of the metabolic interactions between human host and gut microbiomes in health and diseases states.

## Introduction

Primary and secondary bile acids play an important role in human health. As microbial dysbiosis can alter the bile acid pool, it affects human health. The human liver synthesizes two primary bile acids, cholate (CA) and chenodeoxycholate (CDCA), which are conjugated with glycine or taurine and excreted into bile [1]. Their function is to make dietary lipids more soluble. The microbiota in the human gut biotransforms primary bile acids into a variety of secondary bile acids, with the lithocholate (LCA) and deoxycholate (DCA) being most abundant [2]. This microbial transformation results in the bile acid pool becoming more hydrophobic, which is associated with higher toxicity for both human and microbial cells [3]. The corresponding microbial genes for bile acid transformation have been identified [2].

A link between microbial bile acid metabolism and inflammatory bowel disease (IBD), i.e., ulcerative colitis and Crohn’s Disease, has been repeatedly demonstrated [4]. In IBD patients, fecal conjugated bile acid levels are higher while secondary bile acid levels are lower, and the deconjugation and transformation abilities of IBD-associated microbiomes are impaired [4]. Previously, a MetaHIT dataset, including 21 ulcerative colitis (UC) patients, four Crohn’s disease (CD) patients, and 14 healthy controls [5], was analyzed *in silico* revealing a reduced abundance of the bile salt hydrolase (*bsh*) gene in Firmicutes in the four CD patients and an increased abundance of the *bsh* gene in the Actinobacteria phylum in UC patients [6]. In another study [7], a significant reduction of the Firmicutes *bsh* gene was found in UC but not CD patients. Moreover, a reduction of the hydroxysteroid dehydrogenase (*hsdH*) gene in UC patients as well as an increase of the same gene in CD patients compared with controls has been reported [7]. Other diseases that have been associated with alterations of the intestinal bile acids pool include liver cirrhosis, liver cancer, irritable bowel syndrome, short bowel syndrome, and obesity [8–10]. In contrast, the secondary bile acid ursodeoxycholate (UDCA), and its taurine and glycine conjugates, are cyto-and neuroprotective. They have been proposed as potential treatment for neurodegenerative diseases, such as Alzheimer’s and Parkinson’s Disease [11], as well as for IBD [12]. Moreover, bile acids have antimicrobial effects both directly due to their hydrophobicity and indirectly via their stimulation of farnesoid X receptor (FXR), which results in production of antimicrobial peptides [8]. For example, LCA and DCA produced by *Clostridium scindens* have been shown to inhibit the pathogen *Clostridium difficile* in a dose-dependent manner [13].

Bile acid levels are specific to an individual due to individual gut microbiome compositions. Consequently, alterations in bile acid human-microbiome co-metabolism in disease states are also individual-specific and interventions should be targeted. By taking the individual-specific composition of the bile acid pool into consideration, interventions resulting in more favorable bile acid levels could be proposed [2]. Computational systems biology is emerging as a valuable method for predicting individual-specific dietary and drug interventions, which could improve treatment efficacy and reduce the disease burden [14].

One well-established computational approach for modeling human and microbial metabolic states is Constraint-based Reconstruction and Analysis (COBRA). The COBRA approach relies on genome-scale reconstructions of the target organism that represent the metabolism of the target organism in a structured manner and have been manually curated starting from the genome sequence against the available genomic data and literature following established protocols [15, 16]. Genome-scale reconstructions can readily be converted into mathematical models [15]. By enforcing constraints, e.g., nutrient uptake, on the model, condition-specific biological properties of the target organism then can be computed [15]. A common method to compute functional states is flux balance analysis (FBA) [17], in which a biological objective, e.g., biomass production, is optimized. The optimal solution provides a flux value for each model reaction that is consistent with the applied constraints and supports the optimal objective value. Typically, there are alternate optimal flux distributions that result in optimal states [15]. The extent to which each reaction can vary in its flux value, while still being consistent with the applied constraints and providing support for the optimal objective value, can be determined through flux variability analysis (FVA) [18]. The COBRA approach has been successfully applied to model human metabolism in disease states [19]. Moreover, COBRA has valuable applications for interrogating the metabolic capabilities of human intestinal microbes [20]. For instance, using the COBRA approach, the metabolic cross-feeding in small-scale communities of human gut microbes was predicted [21–23]. An advantage of the COBRA approach is that the underlying genome-scale metabolic networks enable mechanistic predictions of metabolic fluxes [15] such that the contribution of each individual microbe in the community can be exactly quantified. Unlike calculating reaction abundances from metagenomic data, this method also takes into account such biological features as substrate availability or species-species boundaries. In the largest-scale gut microbiome modeling study so far, a model community of 11 gut microbes was integrated into a computational framework with the human reconstruction Recon2 [24] and the gut microbial contributions to health-relevant human metabolites were computed [25].

Recently, a comprehensive collection of curated genome-scale reconstructions for 773 human gut microbial strains, AGORA, has been published [26]. With AGORA covering 13 phyla commonly found in human gut microbiomes [26], it becomes possible to construct constraint-based community models that are representative of the human gut microbiome’s composition. AGORA has been curated for a number of gut-specific subsystems, including fermentation, carbon source biosynthesis, respiration, and vitamin biosynthesis [26]. However, AGORA does not account for bile acid transformations. The present study fills this gap.

Based on a comparative genomic approach on the strain level, the metabolic reconstructions in the AGORA compendium were updated to include a functional subsystem for bile acid deconjugation and transformation reactions and are freely available on the VMH website [27]. By expanding the previously developed computational framework [25] both in scale and functionality, we constructed community models on the whole microbiome scale from publicly available metagenomic data. Subsequently, we quantified the individual-specific bile acid production potential in microbiomes from healthy individuals and IBD patients. This mechanistic, microbiome-wide modeling approach can be readily combined with a genome-scale model of human metabolism [24] and customized with individual-specific diets enabling personalized modeling in health and disease states.

## Results

In order to investigate the microbial bile acid production capabilities in individual microbiomes of healthy and IBD individuals, we employed state-of-the-art genomic and computational modeling methods. First, we performed a comparative genomic analysis of bile acid pathways on 693 gut microbial genomes and expanded the microbial metabolic reconstructions present in AGORA accordingly. We then joined the AGORA models with bile acid production pathways into pairwise models and predicted the potential to cooperatively produce secondary bile acids. While each microbe could only produce a subset of the 13 secondary bile acids *in silico*, microbial pairs could produce up to 12 of the 13 bile acids, which highlights that bile acid biotransformation is a microbial community task. Subsequently, we constructed functional and individual-specific gut microbiome models from AGORA using metagenomics data from healthy and IBD individuals to predict the individual bile acid biosynthesis potential. We found inter-individual variation in the production capability of bile acids in healthy individuals as well as significant differences between healthy and IBD microbiomes. Moreover, we were able to compute the contribution of each strain to bile acid deconjugation and transformation while taking the metabolic network of the whole microbiome community and the applied constraints (e.g., dietary uptake) into account.

### Presence of bile acid biotransformation genes in 693 analyzed genomes

We performed a systematic comparative genomic analysis of the bile acid deconjugation and transformation pathway in 632 AGORA genomes, 38 genomes of newly reconstructed organisms, and 23 additional genomes (693 analyzed genomes total) that were available for analysis in the PubSeed database (Figure 1a). The bile salt hydrolase gene (bsh), which encodes the deconjugation of conjugated primary bile acids, was found in 204 of 693 analyzed genomes, including two archaeal genomes, *Methanobrevibacter smithii* ATCC 35061 and *Methanosphaera stadtmanae* DSM 3091 (Table S1). The distribution of the *bsh* gene in Actinobacteria, Bacteroidetes, Firmicutes, as well as the two archaea (Figure 2, Figure S1-2) was in line with previously reported results [28]. Additionally, the *bsh* gene was found in 22 Proteobacteria genomes (Table S1). Among all analyzed hydroxysteroid dehydrogenases, 7a-HSDH was the most widespread enzyme as it was found in 36 of 693 analyzed genomes (Table S1). Additionally, 3α-, 3β-, and 7β- hydroxysteroid dehydrogenases (HSDHs) were found in 14, seven, and three genomes, respectively (Table S1). The *bai* genes for the multistep 7α/β-dehydroxylation pathway were found in seven analyzed genomes belonging to *Clostridioides* sp., *Lachnoclostridium* sp., and *Eggerthella* sp. (Figure S3). Remarkably, all these genomes also have genes for 7α-HSDH or genes for both 3α- and 3β- HSDHs (Table S1). Thus, these organisms could play a crucial role in biotransformation of bile acids in human intestine. At the end, 238 of 693 analyzed organisms were found to be capable of bile acid deconjugations and biotransformation, including 217 reconstructed AGORA organisms (Figure 1a). The corresponding 217 metabolic reconstructions were expanded with the appropriate metabolites and reactions following established procedures [16] (see Methods). The complete reconstructed bile acid biotransformation subsystem contains 38 bile acid metabolites and 82 reactions (Figure 2, Table S2a-b). For CA, CDCA, and the 13 secondary bile acids, transport reactions enabling the uptake and secretion of these metabolites were also added to the corresponding reconstructions.

**Figure 1:**
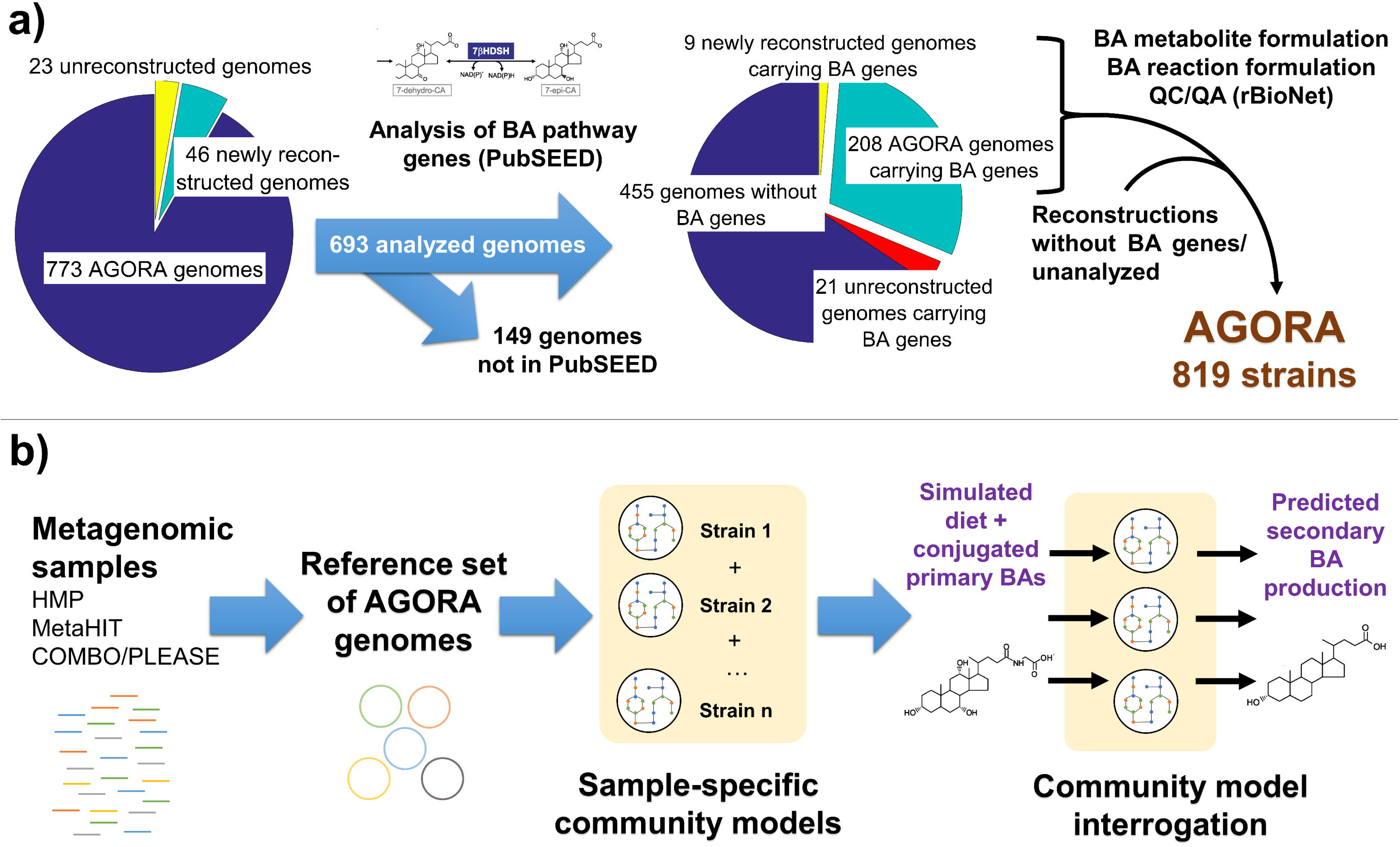
Schematic overview of the workflow in this study. a) Comparative genomic and metabolic reconstruction approach used to expand the AGORA [26] resource with a bile acid (BA) deconjugation and biotransformation subsystem. The comparative genomic approach was performed in the PubSEED [44, 45] platform. Quality control/quality assurance (QC/QA) during reaction and metabolite formulation and addition to the AGORA reconstructions was ensured by using the reconstruction tool rBioNet [59]. b) Computational pipeline used to predict the sample-specific bile acid deconjugation and biotransformation by human gut microbiomes. First, publicly available metagenomic data was retrieved from HMP [29], MetaHIT [5], and the COMBO/PLEASE [32, 61] cohort. Next, the strain-level abundances were mapped onto the reference set of AGORA genomes. Microbial community models were constructed using the illustrated workflow and they account for the strain-level composition of each individual microbiome. Finally, each community model was contextualized with an “Average European” diet supplemented with conjugated primary bile acids and its individual-specific primary bile acid deconjugation and biotransformation potential was computed using flux balance analysis [17] and flux variability analysis [18].

**Figure 2:**
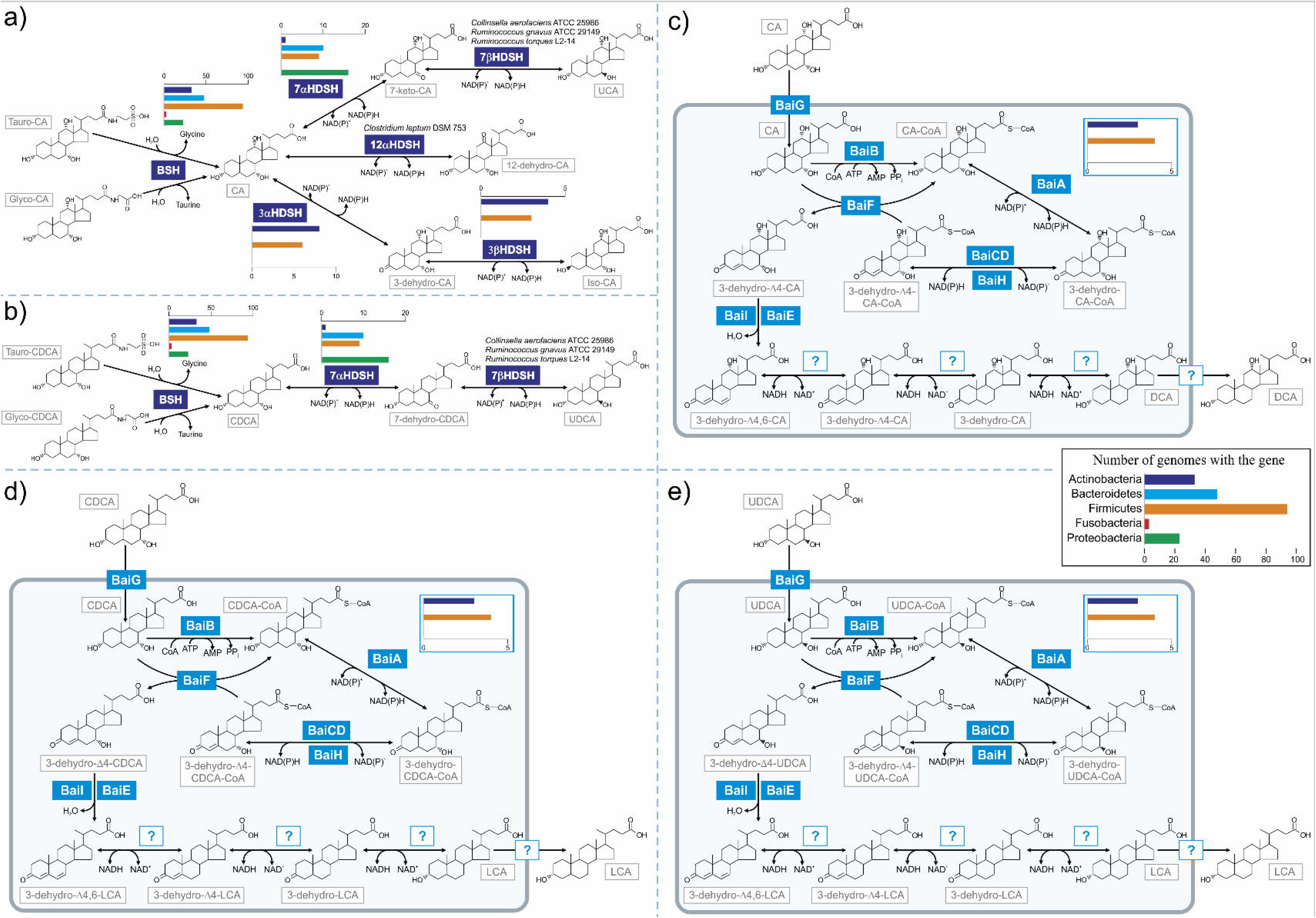
Illustration of bile acid pathways in human gut microbes reconstructed for AGORA. a) Deconjugation of Tauro-CA/Glyco-CA and subsequent conversion to 12-dehydro-CA, UCA, and Iso-CA. b) Deconjugation of Tauro-CDCA/Glyco-CDCA and subsequent conversion to UDCA. c) Conversion of CA to DCA via the *bai* pathway. d) Conversion of CDCA to LCA via the *bai* pathway. e) Conversion of UDCA to LCA via the *bai* pathway. CoA: Coenzyme A. For metabolite abbreviations see Table 1.

### Bile acid transformation capabilities are complementary

The majority of primary bile acids, released by the human gallbladder into the intestine, where the gut microbiome encounters them, are conjugated to glycine or taurine [1]. However, many strains capable of synthesizing secondary bile acids do not possess the bile salt hydrolase (Table S1). To which extent can the 217 bile acid subsystem-carrying strains use the conjugated bile acids as substrates? In order to answer this question, the 217 AGORA reconstructions were converted into condition-specific models by applying an Average European (AE) diet supplemented with taurocholate (Tauro-CA), glycocholate (Glyco-CA), taurochenodeoxycholate (Tauro-CDCA), and glycochenodeoxycholate (Glyco-CDCA) (Table S3) as modeling constraints. The theoretical production potential for each CA, CDCA, and the 13 secondary bile acids (Table 1) was predicted (see Methods). As expected, all 185 reconstructed strains having the bile salt hydrolase could liberate CA and CDCA from the respective conjugated primary bile acids (Table S3). In contrast, no single strain was capable of synthesizing 12-dehydrocholate (12-dehydro-CA), ursocholate (UCA), and UDCA from the conjugated primary bile acids. Of the five strains carrying the *bai* gene cluster, only *Clostridium hiranonis* TO-931 could synthesize LCA and DCA from the conjugated primary bile acids (Table S4). The observed low number of strains capable of both primary bile acid deconjugation and transformation to secondary bile acids suggests the possibility of mutualism between species with complementary bile acid conversion capabilities.

**Table 1:** Overview of primary and secondary bile acids. VMH: Virtual Metabolic Human [27].

| Name | Abbreviation | VMH ID | Type | Producer |
| --- | --- | --- | --- | --- |
| Taurocholate | Tauro-CA | tchola | Primary/<br>conjugated | Human |
| Glycocholate | Glyco-CA | gchola |  |  |
| Taurochenodeoxycholate | Tauro-CDCA | tdchola |  |  |
| Glycochenodeoxycholate | Glyco-CDCA | dgchol |  |  |
| Cholate | CA | cholate | Primary/<br>unconjugated | - Human<br>- Released by 185 AGORA strains |
| Chenodeoxycholate | CDCA | C02528 |  |  |
| 12-dehydrocholate | 12-dehydro-CA | 12dhchol | Secondary | *Clostridium leptum* DSM 753 |
| 7-ketodeoxycholate | 7-keto-CA | 7ocholate |  | 31 AGORA strains |
| 7-dehydrochenodeoxycholate | 7-dehydro-CDCA | 7dhcdchol |  |  |
| 3-dehydrocholate | 3-dehydro-CA | 3dhchol |  | 13 AGORA strains |
| 3-dehydrochenodeoxycholate | 3-dehydro-CDCA | 3dhcdchol |  |  |
| Isocholate | Iso-CA | isochol |  | Six AGORA strains |
| Isochenodeoxycholate | Iso-CDCA | icdchol |  |  |
| Lithocholate | LCA | HC02191 |  | Five AGORA strains |
| Deoxycholate | DCA | dchac |  |  |
| Allolithocholate | allo-LCA | alchac |  |  |
| Allodeoxycholate | allo-DCA | adchac |  |  |
| Ursocholate | UCA | uchol |  | Three AGORA strains |
| Ursodeoxycholate | UDCA | HC02194 |  |  |

In order to investigate the cooperative potential in bile acid metabolism, the 217 bile acid-producing AGORA models were joined with every combination thereof, which resulted in 23,653 pairwise models. The production capabilities for the 13 secondary bile acids of the single strains and the pairwise models was compared on the bile acid-supplemented AE diet. We identified over 15,000 cases, in which a microbe complemented the other’s pathways to synthesize a secondary bile acid, whereas the individual strains were incapable to do so (Figure 3a, Table S5). For example, in the pairs of *Clostridium leptum* DSM 753 and any of the 185 bile salt hydrolase carriers, 12-dehydro-CA synthesis from Glyco-CA or Tauro-CA was enabled (Table S5). Moreover, 717 pairs (3%) enabled the synthesis of DCA/LCA and 69 pairs (0.3%) enabled the synthesis of UCA/UDCA from the conjugated primary bile acids (Table S5), which demonstrates that different pairs of microbes result in the combination of different capabilities in the pathway. There was no pairwise combination enabling synthesis of all secondary bile acids as the maximal number of secondary bile acids to be synthesized by any pair was 12 of the 13 (Figure 3a), which suggests that the presence of complementary microbes could be a bottleneck for bile acid biosynthesis. Taken together, these results demonstrate that bile acid biotransformation is a microbial community task and that the synthesis requires specific strain-strain combinations.

**Figure 3:**
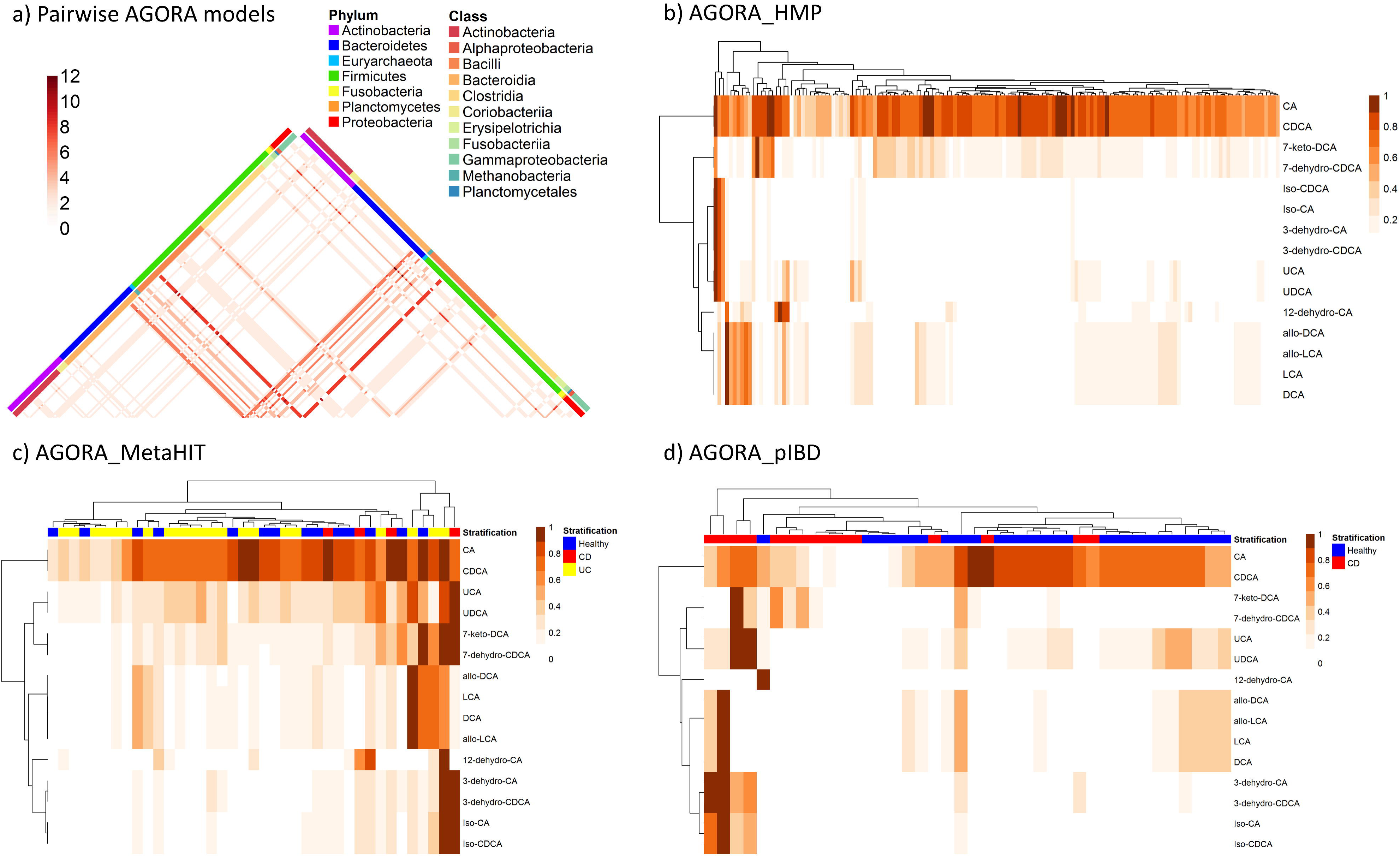
The predicted bile acid production potential of individual gut microbiomes. a) Complementary bile acid biosynthesis capabilities of the 211 AGORA strain models joined in all possible combinations. The numbers of secondary bile acids (out of 13), which can be produced by each pair, are shown. b-d) Bile acid production potential in the three sets of community models constructed in this study, normalized by the highest production (mmol.g_Dw_^−1.^hr^−1^) achieved in any model per bile acid. b) AGORA_HMP, c) AGORA_MetaHIT (10% genome coverage cutoff), d) AGORA_pIBD (10% genome coverage cutoff). AGORA_HMP: models constructed from HMP [29] data, AGORA_MetaHIT: models constructed from MetaHIT [5] data, AGORA_pIBD: models constructed from COMBO/PLEASE [32, 61] data, UC: Ulcerative colitis individuals (AGORA_MetaHIT), CD: Crohn’s Disease individuals (AGORA_MetaHIT and AGORA_pIBD). For metabolite abbreviations see Table 1.

### The bile acid deconjugation and transformation potential varies in healthy adults

Microbes in the human intestine do not exist in isolation but have complex metabolic interactions with each other, which, taken together, influence the health of the host. To predict the combined metabolic potential of an individual’s gut microbiome community (Figure 1b), we generated personalized microbiome community models. First, we mapped the strains identified in each sample to the AGORA compendium and combined all corresponding metabolic reconstructions into one community model. Then, we added a community biomass reaction to the model, which consists of the biomass reaction of each species in the sample and for which the stoichiometric coefficient of each strain was set to its relative abundance in the sample. We predicted the theoretical bile acid production potential of each personalized microbiome community model.

We used 149 metagenomic data from healthy American donors aged 18-40 years provided by the Human Microbiome Project Consortium [29] (see Methods) to assess the extent of the bile acid production potential variation in the gut microbiomes of healthy adults. For the corresponding individual 149 microbiome community models (referred to as AGORA_HMP hereafter), the production potential for the two deconjugated primary and 13 secondary bile acids (Table 1) was predicted on the AE diet supplemented with conjugated primary bile acids (Figure 3b). Although all 149 AGORA_HMP models could produce all analyzed bile acids (Table S6), the quantitative production potential varied significantly between the models (Figure 3b). Only a few AGORA_HMP models produced high levels of LCA and DCA, with differences in quantitative production by a factor of 100 (Figure 3b, Table S6). Microbiomes with low CA/CDCA liberation potential from the conjugated bile acids also had a low secondary bile acid potential, which confirms that the bile salt hydrolase is the gateway reaction in the pathway [30].

In order to determine the overall bile acid production potential, we used flux variability analysis [18, 31] (see Methods). The difference between the maximally and minimally possible flux values (i.e., the flux span) for each bile acid exchange reaction of each strain to the overall microbial bile acid production potential was found to vary considerable across the 149 AGORA_HMP models (Figure S4). The strains contributing to bile acid deconjugation (resulting in CA and CDCA) could be divided into two groups (Figure S4). The strains in group 1 mostly belonged to the Bacteroidetes and Firmicutes phyla and contributed to bile acid deconjugation in most AGORA_HMP models. The strains in group 2 contributed to overall production in less than 100 models and mainly consisted of Actinobacteria, Fusobacteria, Methanobacteria, and Gammaproteobacteria representatives (Figure S4). Similarly, for 7-ketodeoxycholate (7-keto-DCA)/7-dehydrochenodeoxycholate (7-dehydro-CDCA) production by 7-α hydroxysteroid dehydrogenase, Bacteroidetes and Firmicutes representatives contributed to overall production in most AGORA_HMP models, while Gammaproteobacteria strains contributed in only few (Figure S4). Taken together, we predicted inter-person variability in the bile acid biosynthesis potential in healthy individuals, depending on the presence and relative abundance of individual strains in the microbiome samples. Our results suggest that the individual risk of secondary bile acid-induced toxicity would also vary.

### IBD-associated microbiomes are depleted in bile acid deconjugation and transformation capability

In order to explain the reported changed bile acid metabolism in individuals with CD and UC[4], we used strain-level relative abundance data from metagenomics samples from 39 Spanish individuals (21 UC patients, four CD patients, and 14 healthy controls) provided by MetaHIT [5] to construct individual-specific microbiome models, referred to as AGORA_MetaHit hereafter. We then computed the bile acid production potential of each personalized AGORA_MetaHIT model under bile acid supplemented AE diet (Figure 3c-d, Table S7-8) as well as the flux spans for each strain-specific exchange reaction (Figure S5). A statistical analysis (Wilcoxon rank sum test) was performed on total community production potential (mmol*g_Dw_^−1^*hr^−1^), total reaction abundance, reaction abundance on the genus level, and flux spans on the strain level in order to identify the features causing the separation between IBD patients and healthy controls on a more detailed taxonomical level (Table S9-10). In total 40 features differed significantly (p-value <0.05) between the UC patients and healthy controls in AGORA_MetaHIT (Table S9). The quantitative CA and CDCA production (reflecting bile salt hydrolase activity) was significantly lower in UC patients compared with the healthy controls (p-value 0.027, Table S9). In agreement with the findings of Labbe et al [7], the lower production potential could be explained by a significantly lower total abundance of the BSH reaction in the UC individuals (p-value 0.030, Table S9). On the genus level, the BSH reaction abundance was reduced in the *Anaerostipes* (p-value 0.038) and *Coprobacillus* (p-value 0.037) genera in the Firmicutes phylum (Table S9). Moreover, the flux spans for 24 exchange reactions differed significantly between healthy individual and UC patients. For example, UC patients had fewer contributions of *Coprobacillus* sp. 29_1 to deconjugation (p-value 0.040) but more contributions of *Bacteroides fragilis* YCH46 (p-value 0.010) (Table S9).

Due to the small number of CD individuals in the MetaHIT dataset, we used a second metagenomic dataset to build personalized microbiome models (AGORA_pIBD), consisting of 15 children with newly diagnosed CD and of 25 healthy controls [32]. We then repeated the simulation for bile acid production as well as the flux variability analysis on the microbial exchange reactions. Using a principal coordinate analysis on the flux spans, we observed a clear separation into pediatric CD and control models (Figure 4a, b). The CD patients were consistently depleted in contributions of strains to bile acid deconjugation and transformation. This was especially the case for the secretion of CA and CDCA by representatives of the Bacteroidia class (Figure 4a), in line with the report that the dysbiotic pediatric CD patients have lower abundance of the *Bacteroides* genus [32]. In contrast, representatives of the *Escherichia* genus in the Gammaproteobacteria class contributed to bile acid deconjugation and 7-keto-DCA/7-dehydro-CDCA production mostly in the CD microbiomes (Figure 4a), which is consistent with the fact that CD microbiomes are enriched in *Escherichia* sp. [32].

**Figure 4:**
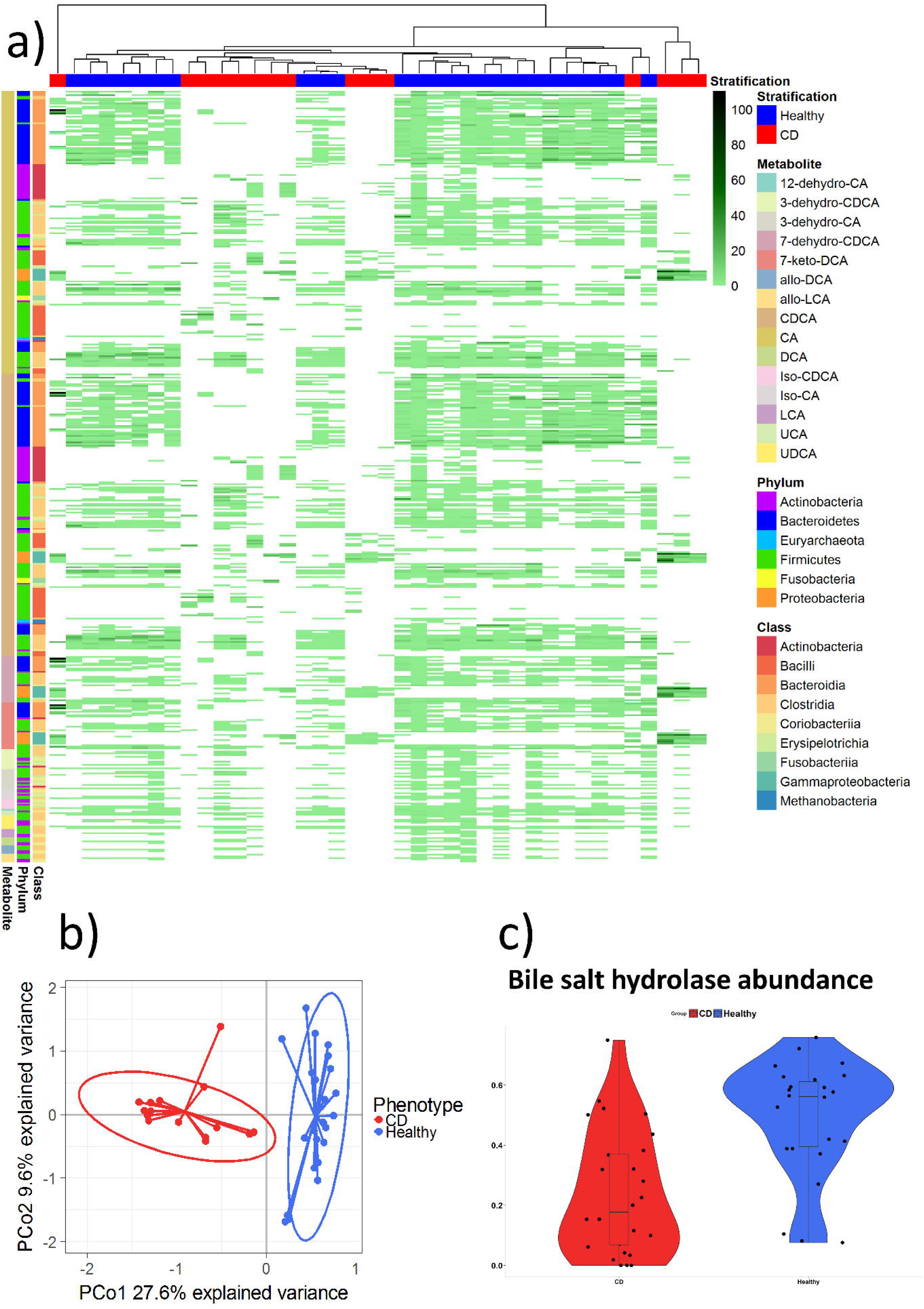
Features of AGORA_pIBD. a) Flux spans for 40 individual models (10% genome coverage cutoff) on all strain to lumen exchanges that carried flux for bile acids in at least one model. For metabolite abbreviations see Table 1. b) Principal Components Analysis (PCoA) of the flux spans depicted in 4a). c) Total abundance of the bile salt hydrolase reaction in the 40 individual models in AGORA_pIBD.

Performing the statistical analysis (Wilcoxon rank sum test) on total community production potential (mmol*g_Dw_^−1^*hr^−1^), total reaction abundance, reaction abundance on the genus level, and flux spans on the strain level revealed that 462 features differed significantly between the pediatric CD and control models (p-value <0.05, Table S10). The CD models had a significantly lower potential to liberate CA and CDCA by deconjugation compared with the control models (p-value 0.013, Table S10), and were also depleted in potential to synthesize secondary bile acids, e.g., 12-dehydro-CA (p-value <0.001, Table S10). These differences could be explained by a lower abundance of most reactions in the bile acid pathway in the pediatric CD individuals on the total community level, e.g., BSH (p-value <0.001, Figure 4c, Table S10). On the genus level, BSH abundance in CD microbiomes was lower in several representative genera of the Bacteroidetes (e.g., *Alistipes, Bacteroides)* and the Firmicutes phyla (e.g., *Coprococcus, Eubacterium, Roseburia, Ruminococcus,* and *Ruminiclostridium*) (p-values for all <0.001, Table S10). In contrast, the CD models had a significantly higher abundance of the BSH reactions in the genera *Blautia* (p-value 0.025), *Enterococcus* (p-value 0.003), and *Escherichia* (p-value 0.002) (Table S10). Several of these genera were also found to separate the pediatric CD patients from controls in the original study [32]. The abundances of the bile acid transformation reactions on the genus level also differed significantly between CD models and controls. The CD models had significantly lower abundance of hydroxysteroid dehydrogenase reactions in several genera, e.g., *Collinsella, Holdemania,* and *Ruminiclostridium* (p-value for all <0.001, Table S10), but higher abundance in the *Escherichia* genus (p-value 0.002) (Table S10). There were also significant differences between CD models and control models in the strain-specific contributions to bile acid production, confirmed by the flux spans of 118 exchange reactions (Table S10). Strain contributions most depleted in the CD models included those of *Bacteroides vulgatus* ATCC 8482, *Ruminococcus (Blautia) torques* L2-14, *Ruminococcus* sp. SR1-5, *Eubacterium ventriosum* ATCC 27560, and *Eubacterium rectale* ATCC 33656 (p-value for all <0.001, Table S10). In contrast, the CD models were highly enriched by contributions of the pathogenic *Escherichia coli* strains O157-H7 Sakai and UTI89 UPEC to deconjugation and 7-keto-DCA/7-dehydro-CDCA production (p-value <0.001, Table S10).

Taken together, both UC and pediatric IBD microbiomes were depleted in bile salt hydrolase activity of *Bacteroides* sp., *Coprobacillus* sp, and *Parabacteroides* sp. (Table S9-10). Compared to UC microbiomes, pediatric IBD microbiomes showed larger differences to healthy controls, as they were additionally depleted in contributions of a variety of other commensal microbes to bile salt hydrolase and to bile acid transformation but enriched in contributions of pathogenic *Escherichia sp*. (Table S10). Thus, the pediatric IBD microbiomes clearly differed in bile acid deconjugation and transformation potential while the differences between UC microbiomes and healthy were less pronounced and limited to bile acid deconjugation potential.

### Complementary microbe-microbe interactions are required for secondary bile acid biosynthesis in individual communities

Can the bile acid biotransformation potential of a community be predicted from strain and gene abundance alone? To test whether the bile acid production potential can be directly inferred from the abundance of the synthesizing reaction, we calculated the Pearson correlation between the individual production potential for the two deconjugated primary and 13 secondary bile acids (Table 1) and the total community abundance for all reactions in the bile acid pathway in AGORA_HMP, AGORA_MetaHIT, and AGORA_pIBD (228 community models in total). For 13 of the analyzed bile acids, the correlation between production potential and the abundance of reaction directly synthesizing the respective bile acid was 0.93 or higher, as expected (Table S11). In contrast, the Pearson correlation between production potential for UCA and UDCA and the abundance of the 7-β hydroxysteroid dehydrogenase reaction (VMH ID: UCA7bHSDHe/UDCA7bHSDHe) calculated for all 228 models was 0.78 indicating that high reaction abundance does not always correspond to high production. Plotting the total reaction abundance of the 7-β hydroxysteroid dehydrogenase yielding UDCA (UDCA7bHSDHe) against the UDCA biosynthesis potential in all 228 models revealed that in 22 models, the total production potential was lower than one would expect from the total 7-β hydroxysteroid dehydrogenase reaction abundance (Figure 5c). For instance, the two models with the highest reaction abundance (both of them of pediatric CD microbiomes) only achieved a low UDCA production flux (Figure 5c). This result shows that factors other than the abundance of strains with 7-β hydroxysteroid dehydrogenase activity limited the production flux. Thus, the bile acid production potential cannot be predicted from species and gene abundances alone in all cases and metabolic fluxes in the community should be considered. Constraint-based modeling is ideal for such analyses of metabolic dependencies since it is mechanistic, quantitative, and takes species-species metabolic exchanges and boundaries into account [33].

**Figure 5:**
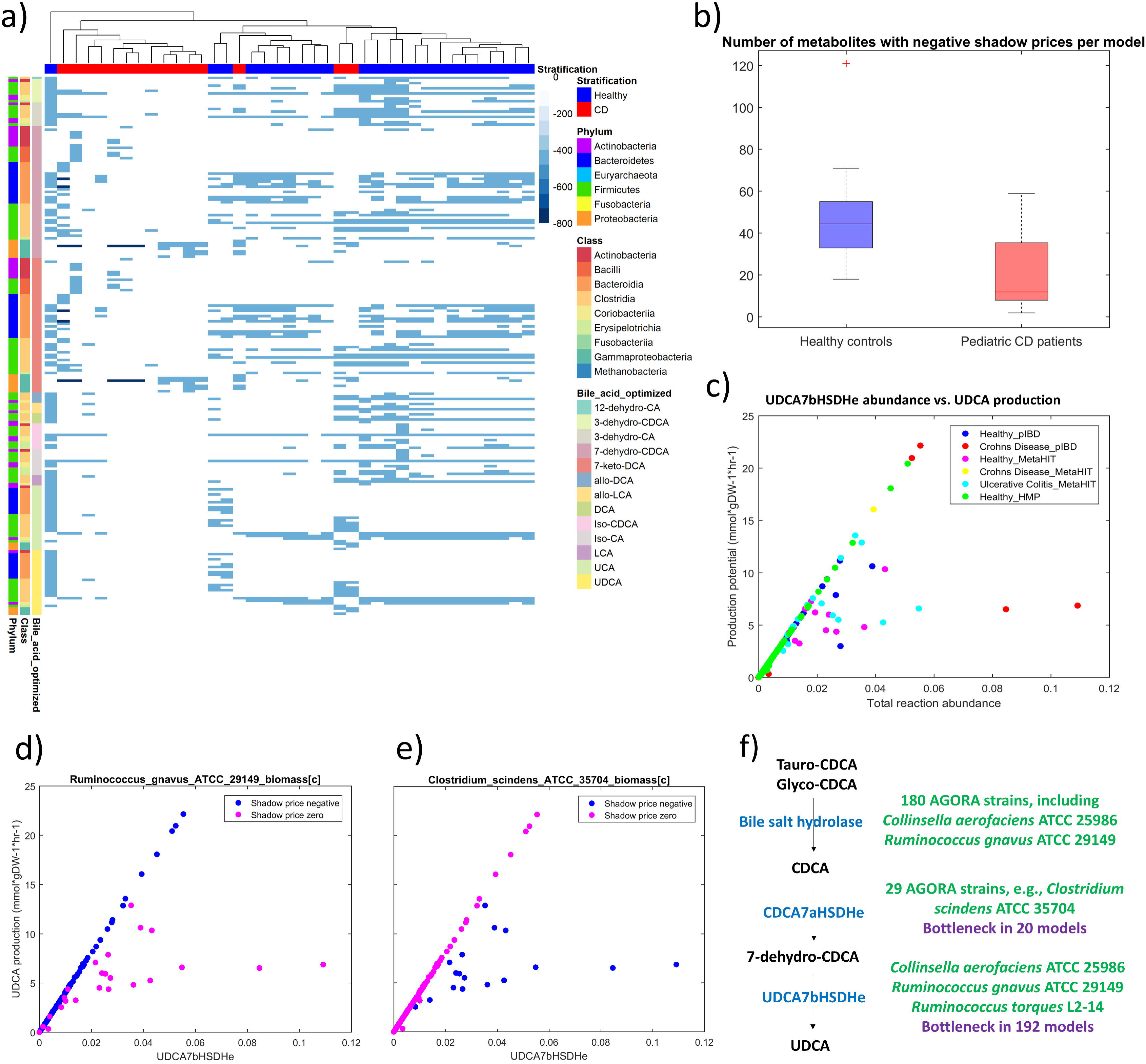
Depiction of shadow prices computed when optimizing production of total community secondary bile acid production. Positive shadow price indicates that increasing availability of the metabolite would increase bile acid biosynthesis. a) Shadow prices in AGORA_pIBD. The columns represent individual communities (blue=healthy controls, red=pediatric CD microbiomes) and the columns represent all metabolites that had a high negative shadow price (£-400) in at least one community model. The metabolites are annotated by taxonomy. The ‘bile acid optimized’ color bar shows the bile acid for which production was optimized. b) Number of metabolites with negative shadow prices in the pediatric CD microbiome models and healthy controls models. c) Total abundance of the UDCA7bHSDHe reaction plotted against UDCA production potential in all 228 microbiome models. d) Shadow prices for the biomass metabolite of *Ruminococcus gnavus* ATCC 29149, an UDCA7bHSDHe carrier, plotted against UDCA production and UDCA7bHSDHe abundance in the same models as in 5c). A negative shadow price shows that the abundance of *R. gnavus* is limiting UDCA production. e) Shadow prices for the biomass metabolite of *Clostridium scindens* ATCC 35704, a CDCA7aHSDHe carrier, plotted against UDCA production and UDCA7bHSDHe abundance in the same models as in 5c). A negative shadow price shows that the abundance of *C. scindens* is limiting UDCA production. f) Schematic description of the UDCA biosynthesis pathway (more detail in Figure 2b). Steps that are bottlenecks for UDCA production in certain community models, indicated by a high negative shadow price for biomass metabolites of strains carrying the respective reaction, are shown. UDCA7bHSDHe: VMH ID for 7-beta-hydroxysteroid dehydrogenase, UDCA7bHSDHe: VMH ID for 7-beta-hydroxysteroid dehydrogenase, VMH: Virtual Metabolic Human [27]. For metabolite abbreviations see Table 1.

What are the factors in the metabolic network that are limiting the entire community’s production potential for secondary bile acids? This question can be answered by investigating the shadow prices of the flux balance analysis solution. The shadow price is a measurement for the value of a metabolite towards the optimized objective function, with a negative shadow price indicating that the flux through the objective function would increase when the availability of this metabolite would increase [34]. In order to identify limiting factors for secondary bile acid production, we investigated the shadow prices in the flux balance analysis solutions (see Methods) computed for all three sets of models when optimizing production of the 13 secondary bile acids (Figure 5a, Figure S6-7, Table S12). Large negative shadow prices (value ≤-400) were found for biomass metabolites of 108 strains carrying bile acid enzymes (Table S12). Thus, the abundance of these strains was limiting secondary bile acid production in at least one microbiome model. Note that only biomass metabolites in strains already present in the individual (and thus in the model) can have shadow prices. When comparing pediatric CD patients and controls in the AGORA_pIBD models (Figure 5a), the number of metabolites with negative shadow prices was significantly lower in the CD models than in control models (19.9 +/- 18.0 vs. 49.2 +/- 24.2, p-value 0.0002, Wilcoxon rank-sum test) (Figure 5b). Hence, due to the known dysbiosis in the pediatric CD patients [32], they were depleted in strains with bile acid biosynthesis capabilities. This results highlights that an increase in secondary bile acid biosynthesis in these individual communities could only be achieved by introducing additional microbial strains.

The shadow prices for UDCA biosynthesis (Table S12) were inspected in all 228 models to identify limiting factors (Figure 5d-f). Note that shadow prices for UCA are analogous. Of the reconstructed strains, only *Collinsella aerofaciens* ATCC 25986, *Ruminococcus gnavus* ATCC 29149, and *Ruminococcus torques* L2-14 carried the 7-β hydroxysteroid dehydrogenase and were thus capable of UDCA biosynthesis (Table S1). In eight microbiomes, all of which were of pediatric CD patients, all three strains were absent in the corresponding metagenomic samples and thus in the models. Consequently, these models were unable to produce any UDCA (Table S8) and all shadow prices for UDCA production were zero (Table S12). In 198 models, shadow prices were negative for the biomass metabolites of one or more of the three UDCA-producing strains, as expected (Table S12). In contrast, in 22 microbiome models, shadow prices were zero for the three strains even though UDCA was produced (Table S12). Instead, shadow prices were negative for the biomass metabolites of one or more of 24 other strains including *Bacteroides* sp., *Escherichia* sp., Clostridiales, and Erysipelotrichales representatives (Table S12). These 22 microbiome models were the same models that had a lower UDCA production potential than expected (Figure 5c). All 24 strains were found in the analysis of complementary pairs to enable UDCA biosynthesis (Table S5) when pairwise joined with *C. aerofaciens,* and *R. gnavus*, and are characterized by the presence of the 7-α hydroxysteroid dehydrogenase yielding 7-dehydro-CDCA (VMH ID: CDCA7aHSDHe) (Figure 5f, see also Figures 2b) and 2e)). Only ten of the 24 strains could enable UDCA biosynthesis when joined with *R. torques* (Table S5). This is due to the fact that *R. torques* does not possess the bile salt hydrolase and thus requires one of the ten strains that carry both the 7-α hydroxysteroid dehydrogenase and the bile salt hydrolase reaction to complement its UDCA biosynthesis pathway (Figure 5f).

Shadow prices were plotted against UDCA7bHSDHe abundance and UDCA production exemplarily for two strains present in many of the 228 microbiomes in order to confirm that the UDCA production in 22 models was limited by the availability of strains possessing the 7-α hydroxysteroid dehydrogenase: (i) *R. gnavus* ATCC 29149 (Figure 5d), and (ii) *Clostridium scindens* ATCC 35704, a strain carrying 7-α hydroxysteroid dehydrogenase (Figure 5e). In microbiome models for which 7-β hydroxysteroid dehydrogenase abundance directly correlated with UDCA production and in which *R. gnavus* was present, shadow prices for *R. gnavus* biomass were negative (Figure 5d). In the 22 models showing lower UDCA production than expected and in which *C. scindens* was present, shadow prices for *C. scindens* biomass were negative (Figure 5e). This example illustrates that the biosynthetic capabilities of a strain need to be viewed in the context of the entire gut microbiome community’s metabolic network while also taking metabolic constraints (e.g., substrate availability) into account. In fact, the correlation between total abundance of the 7-α hydroxysteroid dehydrogenase reaction and UDCA production potential was only 0.05 (Table S11), which makes it non-obvious without inspection of the flux solutions that this reaction was synthesis-limiting in the 22 microbiomes.

In summary, by analyzing the shadow prices in the FBA solutions when optimizing secondary bile acid production, strains that could increase their biosynthesis potential were uniquely identified for each personalized community model. The analysis revealed that in certain models, the abundance of microbes possessing the 7-α hydroxysteroid dehydrogenase reaction, which provides the necessary precursor for UDCA production was synthesis-limiting, resulting in lower UDCA production than one would expect from the abundance of the 7-β hydroxysteroid dehydrogenase. This result highlights once more that bile acid deconjugation and transformation is a cooperative task and needs to be viewed in the context of the community-wide metabolic network. This analysis allows for the generation of mechanistic hypotheses, e.g., bottlenecks that are limiting the transformation of primary to secondary bile acids, and confirms the known fact that IBD patients have altered bile acid profiles [4].

## Discussion

In this work, we performed a comparative genomic analysis of the bile acid deconjugation and transformation pathway (Figure 1a, Figure 2) in 693 human gut microbe genomes. The reactions encoded by the identified genes were reconstructed and added to a resource for genome-scale metabolic reconstructions, AGORA [26]. The resource is publicly available at the VMH website [27]. From AGORA, pairwise models and personalized gut microbiome models (Figure 1b) were built and extensively interrogated for single, pairwise, and community-wide bile acid production potential, as well as flux contributions and flux-limiting factors in individual communities.

We demonstrated that microbes can complement each other’s bile acid pathway (Figure 3a). For example, no single strain, but 69 pairs of bacteria could cooperatively convert tauro-or glyco-CDCA into UDCA. Since UDCA is much more hydrophilic and less toxic to bacteria than CDCA [3], cooperatively performing this task would be of advantage to both microbes in these pairs. Microbe-microbe co-presence were also found to be limiting for UDCA biosynthesis in the microbiome community models (Figure 5c-f) with possible implications for human health since UDCA is a cytoprotective bile acid with therapeutic potential [3, 11]. For example, UDCA was found to improve bile acid homeostasis and suppress ileitis in a model of Crohn’s Disease [12].

While it can be intuitively understood that bile acid biosynthesis is a cooperative task in the gut microbiome from the known fact that no strain possesses the complete pathway [2], with constraint-based modeling, these microbe-microbe metabolic dependencies can be exactly predicted. Each personalized gut microbiome community model constructed from AGORA can be interrogated for the contribution of each microbe to the overall production of a bile acids (Figure 4a, Figure S4-5). The models may also be interrogated for microbe-microbe cross-feeding which is necessary for the synthesis of the end products of the pathway. Due to the underlying genomescale metabolic reconstructions in the community models, mechanistic explanations for predicted synthesis bottlenecks and microbe-microbe metabolic dependencies can be derived from the simulation results.

Using personalized microbiome models constructed from AGORA (Figure 1b), we could predict the variety in bile acid production across healthy individuals (Figure 3b) and identify differences in bile acid production potential between IBD patients and healthy controls (Figure 3c-d). Moreover, the exact contribution of each strain to the predicted overall production was found to be variable across individuals (Figure 4a, Figure S4-5). Constraint-based modeling can thus elucidate the importance of individual microbes towards fulfilling a community task and mechanistically link produced metabolites to species beyond correlations. For example, the correlations between bacterial abundance and gene pathway enrichment has been analyzed in the pediatric Crohn’s Disease dataset for a variety of pathways [32], and for secondary bile acid biosynthesis, a positive correlation with *Bacteroides* sp. has been found. Consistently, we predicted that a variety of *Bacteroides* strains performed bile acid deconjugation and 7-keto-DCA/7-dehydro-CDCA biosynthesis, and their contribution was significantly depleted in the Crohn’s Disease microbiomes (Table S10). A recent study investigated the microbiomes and fecal metabolomes of pediatric IBD patients and their relatives and could distinguish two metabotypes both in patients and relatives [35]. The IBD-associated metabotype was characterized by an altered bile acid profile, with increased levels of cholate and sulfated and taurine-conjugated primary bile acids. This finding suggests a reduced bile acid deconjugation and conversion potential of the gut microbiota [35], which is consistent with our results. Interestingly, the IBD-associated metabotype also had higher levels of 7-keto-CDCA, which may reflect altered secondary bile acid metabolism [35] and could be further investigated in future studies.

While the bile acid pathway is comparatively linear and most steps are only present in a limited number of species, other pathways, such as the citric acid cycle or glutamate metabolism, play a central role in metabolism and are present in a much larger variety of species. The effect of depletion or enrichment of taxa possessing central pathways, such as the citric acid cycle, cannot be easily predicted from gene abundance alone. Top-down multivariate statistical analyses of metabolomic data have identified correlations between microbial species and metabolites [36], however, the underlying mechanisms often remain unexplained. For instance, Lewis et al. found that several pathways, e.g., glycerophospholipid metabolism, amino benzoate degradation, sulfur relay system, and glutathione metabolism, predicted the separation between healthy and dysbiotic microbiomes [32]. It cannot be easily inferred from correlations alone how these differentially enriched pathways would affect the levels of health-relevant metabolites, such as glutathione or phospholipids. Urinary and/or fecal metabolites found to be altered in IBD states include propionate, butyrate, lactate, citric acid cycle intermediates, amino acids, folate, hippurate, formate, taurine, choline, sphingolipids, and polyamines [37–40]. In future efforts, the computational workflow used in this study could be applied to predict the gut microbial metabolome beyond bile acid metabolism in order to gain insights into the mechanisms behind altered metabolomes in disease states.

The gut microbiome community models constructed from AGORA can be readily joined with the human reconstruction Recon3 [41] enabling the computation of human-microbe co-metabolism. For example, we have previously shown that a simplified gut microbe community can produce precursors for human bioactive metabolites including neurotransmitters or glutathione [25]. This approach could be expanded to include the representative microbiome-wide community models constructed in this study. Including the human component of the bile acid subsystem would add another layer of complexity and could reveal unexpected, non-intuitive human-microbe cross-feeding. The possibility that microbes can also produce precursors in the human primary bile acid synthesis pathway may be explored in future efforts. Moreover, reabsorption and detoxification of secondary bile acid in the human liver [42] could be predicted. Recon3 has recently been expanded with a secondary bile detoxification module [41]. For example, investigating the shadow prices for glucuronidated or sulfated LCA or DCA could elucidate bottlenecks in the detoxification of these highly toxic bile acids. Finally, while the present study was simplified by using the same diet and conjugated bile acid uptake constraints in every model, the effect of individual-specific precursor availability and diet could be examined in future studies.

In summary, we constructed for the first time *in silico* personalized gut microbiome models that can elucidate the metabolic potential of an individual’s microbiome. The models can be readily joined with a human reconstruction, e.g., Recon3, and personalized through integration of transcriptomic data [43] or nutritional information via the Virtual Metabolic Human [27] resource. We expect that the AGORA resource supplemented with microbial bile acid metabolism will have valuable applications in unraveling the role of human-gut microbiome metabolic interactions in human health.

## Material and Methods

### Comparative genomic approach

All 773 strains of the AGORA resource [26], 46 new strains with a metabolic reconstruction, and 23 currently not reconstructed strains, were checked for the presence of their genomes at the PubSEED resource [44, 45], resulting in 690 bacterial and three archaeal genomes to be considered in this study (Figure 1a). Note that only 38 of the newly reconstructed microbes had their genome available in PubSEED and were consequently used for the comparative genomic approach. All 693 human gut microbe genomes were analyzed for the presence of orthologs of bile acid deconjugation and biotransformation genes (Table S1). Orthologs are defined as genes that satisfy the following conditions: (1) Orthologs should be closely homologous proteins (e-value cutoff = e-50). (2) Orthologs should be found in the same genomic context, i.e., structure of gene locus should be conserved in related genomes. (3) Orthologs should form a monophyletic branch of a phylogenetic tree.

For the search of homologs and analysis of genomic context, the PubSEED platform was used along with phylogenetic trees for protein domains in MicrobesOnline [46]. Multiple protein alignments were performed using the MUSCLE v. 3.8.31 tool [47, 48]. Phylogenetic trees were constructed using the maximum-likelihood method with the default parameters implemented in PhyML-3.0 [49]. The trees obtained were visualized and midpoint-rooted using the interactive viewer Dendroscope, version 3.2.10, build 19 [50].

The following previously analyzed genes were used as a starting point, (1) genes for bile salt hydrolases (BSH) from multiple genomes [28], (2) 7α-hydroxysteroid dehydrogenase (HSDH) gene from *Bacteroides fragilis* [51], (3) 3α- and 3β-HSDHs genes from *Eggerthella lenta* DSM 2243 and *Ruminococcus gnavus* ATCC 29149 [2], (4) 7α-HSDH and *baiABCDEFGHI* genes for a multistep 7α/β-dehydroxylation pathway, (5) *bai* genes from *Eggerthella lenta* DSM [2], and (6) 7β-HSDH gene from *Clostridium absonum* [52]. BSH proteins are closely related to penicillin V amidase (PVA) ones [28]. To avoid incorrect annotations, a phylogenetic tree for BSH proteins and their homologs in the analyzed genomes was constructed (Figure S1-2) and orthologs of the known BSH genes were identified. All HSDH proteins listed above demonstrated similarity to each other and with BaiA proteins. Thus, orthologs for HSDH/BaiA proteins were resolved with the construction of a phylogenetic tree (Figure S3).

All of the annotated genes are represented as a subsystem at the PubSEED website [53] and can be found in Table S1.

### Formulation and addition of reactions

Reaction mechanisms were retrieved from the KEGG database [54] as well as published literature (e.g., [55]). For all genomes having genes for BSH, HSDHs, or the complete 7α/β-dehydroxylation pathway (Figure 2), metabolic mass- and charge-balanced reactions were formulated. Exchange reactions were added for all extracellular metabolites. The majority of reactions were associated with genes and proteins annotated in the analyzed genomes. Reactions not-associated with genes and proteins were only added if the gene was unknown but the reaction was required to eliminate dead-ends in a metabolic pathway. Thus, the followed gap-filling reactions were added without associations with genes or proteins: (1) The three last steps for the 7α/β-dehydroxylation pathway are still unknown but was predicted to be three NADH-dependent reductases [55]. Hence, these reactions were added without associations to genes and proteins. (2) A transport reaction for LCA, the final product of 7α/β-dehydroxylation pathway was added despite the transporter being currently unknown. Moreover, pathways that yield allolithocholate (allo-LCA) and allodeoxycholate (allo-DCA) were included for strains possessing the *bai* gene cluster as these compounds are known to be side products of the 7α/β- dehydroxylation pathway [56] and found in human adults under certain circumstances [1]. (3) *Clostridium leptum* demonstrates 12α-HSDH activity but the precise gene is not known [57].

Pathways for cholesterol reduction to coprostanol were also reconstructed. These enzymatic activities, both cytoplasmic an extracellular, were demonstrated for *Lactobacillus acidophilus, Lactobacillus bulgaricus*, and *Lactobacillus casei*. Precise mechanisms of these reactions as well enzyme-encoding genes are not known but the biotransformation was shown to be associated with oxidation of NADH to NAD+ [58]. Consequently, reactions for extracellular and cytoplasmic NADH-dependent reduction of cholesterol to coprostanol were added to six *Lactobacillus* sp. models, together with exchange reactions for cholesterol and coprostanol as well as a predicted reaction for cholesterol uptake.

All metabolites and reactions were formulated following an established reconstruction protocol [16]. Metabolites and reaction abbreviations in the bile acid subsystem were created in accordance with the Virtual Metabolic Human (VMH) [27] nomenclature to ensure that they are compatible with the Recon2 [24] and Recon 3 [41] nomenclature. The MATLAB-based reconstruction tool rBioNet [59], which ensures quality control and quality assurance, such as mass- and charge-balance, was used to add the metabolites and reactions to the appropriate reconstructions. All reactions and metabolites in the bile acid subsystem in AGORA are described in Table S2a-b.

### Expansion of AGORA

A total of 46 gut microbial strains were newly reconstructed. The reconstructions were generated by semi-automatically expanding and curating KBase [60] draft reconstructions following the established AGORA pipeline [26] (Table S13). Of the 773 AGORA strains and 46 newly reconstructed strains, 217 strains total carried at least one gene in the bile acid pathway (Table S1) and six produce coprostanol. The corresponding 217 reconstructions were expanded by the appropriate metabolites and reactions (Figure 1a). The expanded resource, accounting for 819 strains, is available on the VMH website [27].

### Construction of pairwise models

The 217 AGORA reconstructions carrying bile acid reactions were joined pairwise in every possible combination as described previously [26] using the createMultipleSpeciesModel function in the COBRA Toolbox [31]. In total, 23,653 pairwise models were created.

### Construction of sample-specific gut microbiota models

Publicly available metagenomic data from 149 individual microbiotas was obtained from the Human Microbiome Project website [29]. The strain-specific abundances in the samples were retrieved as described previously [26], and normalized for each individual to obtain the relative abundances of each strain. Paired-end Illumina raw reads of the Spanish MetaHit cohort, consisting of 21 ulcerative colitis patients, four Crohn’s disease patients, and 14 healthy controls [5], were retrieved from EBI under the accession ERA000116. Paired end Illumina raw reads of 28 dysbiotic Crohn’s disease (CD) patients in the PLEASE cohort [32] and of 26 healthy controls in the COMBO cohort [61] were retrieved from NCBI SRA under SRA: SRP057027. For both datasets, the reads were pre-processed and then mapped onto the reference set of 773 AGORA genomes [62]. In order to reduce the number of false positives, a cutoff of 10% genome coverage was applied to the resulting coverages (representing a threshold of at least 10% genome coverage for each microbe in each human individual). The resulting coverages were normalized for each individual in order to obtain the relative abundances. In order to avoid too small model sizes, microbiome ssamoles, for which less than 20 strains could be mapped to the reference set of AGORA genomes, were excluded from the analysis. This was the case for 13 samples from the PLEASE cohort and one sample from the COMBO cohort.

A microbiome model generation and analysis workflow was developed (Figure 1b). For all strains in the 228 metagenomic samples that could be mapped to AGORA, the corresponding reconstructions were joined together to a constraint-based microbial community model. The resulting sample-specific microbial community models have a compartment that simulates the intestinal lumen, which provides an inlet for the simulated dietary input and allows for microbe-microbe cross-feeding, as previously described [63]. An outlet representing fecal secretion was added. A community-level biomass objective function was formulated from the strain-specific normalized microbial abundances. The lower and upper bounds on the community biomass objective function were set to 0.4 and 1 mmol*g_DW_^−1^*hr^−1^, respectively. Coupling constraints, which enforce the flux through all metabolic and transport reactions to be proportional to an organism’s biomass objective function, were implemented as previously described [63].

### Simulations

All simulations were performed in MATLAB version 2016b (Mathworks, Inc.) using the COBRA Toolbox [31]. Flux balance analysis [17] was performed using the optimization solver CPLEX through the Tomlab (Tomlab, Inc.) interface. For flux variability analysis [18], the fastFVA function was used with the IBM CPLEX (IBM, Inc) solver.

### Definition of the Average European diet

A diet representing the nutrient intake of an average European individual was devised and converted to uptake constraints resources in the Virtual Metabolic Human database (VMH) [27]. The diet was supplemented with metabolites previously determined necessary for the biomass production of at least one AGORA reconstruction [26]. The AE diet was implemented in the AGORA reconstructions as well as the sample-specific community models by constraining the lower bounds on exchange reactions to the defined uptake rate. Note that not every metabolite in the AE diet could be transported by every model. Additionally, in order to enable modeling of bile acid transformation, the uptake of the conjugated primary bile acids Glyco-CA (VMH: gchola), Tauro-CA (VMH: tchola), Glyco-CDCA (VMH: dgchol), and Tauro-CDCA (VMH: tdchola) was allowed by setting the lower bounds on the corresponding exchange reactions to −1000. The lower bounds on all other exchange reactions were set to zero. Constraints applied to simulate the dietary uptake are shown in Table S3.

### Interrogation of models for bile acid synthesis capabilities

The bile acid production potential in 217 AGORA reconstructions, 23,653 pairwise models, and 228 sample-specific community models was computed using FBA [17]. To predict the theoretical bile acid production potential, the exchange reactions (in the single and pairwise models) and the fecal secretion reactions (in the community models) for CD, CDCA, and 13 secondary bile acids were set as the objective function one by one and production flux was maximized. Shadow prices for all metabolites in the respective model were retrieved from the computed FBA solutions [17]. The contributions of individual strains to overall bile acid production were computed as follows: (i) Production flux through the respective the fecal secretion reactions in the community models was maximized. (ii) Flux variability analysis [18] was performed on the luminal exchanges in all strains that could transport the respective bile acid resulting in the minimal and maximal exchange flux values. (iii) For each luminal exchange, the flux span was calculated.

### Data analysis

The calculation of the Pearson correlation and the two-sided Wilcoxon rank-sum test were performed in MATLAB version 2016b (Mathworks, Inc.). All datasets were plotted in MATLAB version 2016b or in R version 3.3.2 [64]. The aheatmap, pheatmap, ggplot2, easyGgplot2, and RColorBrewer packages in R were used for data visualization. Heat maps were generated with the Euclidian distance measure for clustering rows and columns, and complete linkage as the hierarchical clustering method. Principal Components Analysis (PCoA) was performed with the vegan package in R using the Bray-Curtis dissimilarity index.

## Declarations

### Availability of data and materials

The annotated bile acid pathway genes are represented as a subsystem at the PubSEED website [53]. The expanded AGORA reconstructions generated in this study are available at the VMH website [27]. The results of simulations performed in this study are shown in this article and its Supplementary files. Scripts used to analyze simulation results will be available online [65] and rely on the COBRA Toolbox (https://github.com/opencobra/cobratoolbox) [31].

### Author Contributions

AH and IT conceived the study. DAR performed the comparative genomic analysis and formulation of bile acid reactions. AH performed the expansion of AGORA reconstructions, microbiome modeling simulations, and analysis of simulation results. FB retrieved the metagenomics data and constructed the microbiome models. LH and RMTF provided tools and computational infrastructure for large-scale simulations. IT supervised the study. AH and DAR drafted the manuscript. All authors read, edited, and approved the final manuscript.

## Acknowledgements

The authors thank Dr. Eugen Bauer for help with generating the figures and for valuable discussions.

